# Neural Adaptation and Fractional Dynamics as a Window to Underlying Neural Excitability

**DOI:** 10.1101/2022.09.01.506146

**Authors:** Brian N. Lundstrom, Tom Richner

**Author notes:** **Funding:** BNL was funded by NIH NINDS (K23NS112339). **Disclosures:** BNL declares intellectual property licensed to Cadence Neuroscience Inc (contractual rights waived), site investigator (Medtronic EPAS, NeuroPace RESPONSE, Neuroelectrics tDCS for Epilepsy), and industry consultant (Epiminder, Medtronic, Neuropace, Philips Neuro; funds to Mayo Clinic).

## Abstract

The relationship between macroscale electrophysiological recordings and the dynamics of underlying neural activity remains unclear. We have previously shown that low frequency EEG activity (<1 Hz) is decreased at the seizure onset zone (SOZ), while higher frequency activity (1-50 Hz) is increased. These changes result in power spectral densities (PSDs) with flattened slopes near the SOZ, which are assumed to be areas of increased excitability. We wanted to understand possible mechanisms underlying PSD changes in brain regions of increased excitability. We hypothesize that these observations are consistent with changes in adaptation within the neural circuit.

We developed a theoretical framework and tested the effect of adaptation mechanisms, such as spike frequency adaptation and synaptic depression, on excitability and PSDs using filter-based neural mass models and conductance-based models. We compared the contribution of single timescale adaptation and multiple timescale adaptation.

We found that adaptation with multiple timescales alters the PSDs. Multiple timescales of adaptation can approximate fractional dynamics, a form of calculus related to power laws, history dependence, and non-integer order derivatives. Coupled with input changes, these dynamics changed circuit responses in unexpected ways. Increased input without synaptic depression increases broadband power. However, increased input with synaptic depression may decrease power. The effects of adaptation were most pronounced for low frequency activity (< 1Hz). Increased input combined with a loss of adaptation yielded reduced low frequency activity and increased higher frequency activity, consistent with clinical EEG observations from SOZs.

Spike frequency adaptation and synaptic depression, two forms of multiple timescale adaptation, affect low frequency EEG and the slope of PSDs. These neural mechanisms may underlie changes in EEG activity near the SOZ and relate to neural hyperexcitability. Neural adaptation may be evident in macroscale electrophysiological recordings and provide a window to understanding neural circuit excitability.

**Author Summary:** Electrophysiological recordings such as EEG from the human brain often come from many thousands of neurons or more. It can be difficult to relate recorded activity to characteristics of the underlying neurons and neural circuits. Here, we use novel theoretical framework and computational neural models to understand how neural adaptation might be evident in human EEG recordings. Neural adaptation includes mechanisms such as spike frequency adaptation and short-term depression that emphasize stimulus changes and help promote stability. Our results suggest that changes in neural adaptation affect EEG signals, especially at low frequencies. Further, adaptation can lead to changes related to fractional derivatives, a kind of calculus with non-integer orders. Neural adaptation may provide a window into understanding specific aspects of neuron excitability even from EEG recordings.

## Introduction

Neurons are electrically excitable cells that are a fundamental computational unit of the nervous system (1–3). Excitatory and inhibitory neurons interact as neural circuits perform computational tasks. Broadly, excitatory neurons generate action potentials after some stimulus threshold is reached. The rate of action potentials then increases as the stimulus increases, which leads to an increase in downstream excitatory synaptic transmission.

However, multiple negative feedback mechanisms influence excitatory signals. These feedback mechanisms can act to directly decrease the rate of action potentials or can affect synaptic transmission. One example that appears to be present in the vast majority of excitatory neocortical neurons is spike frequency adaptation (SFA) (2,4,5), which is a negative feedback mechanism that slows action potential generation near the cell body at the axon hillock in response to a sustained stimulus (Figure 1). Another example is short-term synaptic depression, a negative feedback mechanism that is a form of synaptic plasticity (6,7). These feedback mechanisms are typically described by a single timescale, such as an exponential time constant, although it is clearly recognized that there are often multiple timescales present. These timescales are generally thought to encompass a range varying from hundreds of milliseconds to several minutes. Over this range, multiple timescales of adaptation can approximate fractional dynamics, a form of power law dynamics with history dependence (8,9). These mechanisms can be considered a form of short-term plasticity and work at a faster timescale than long-term depression and long-term facilitation, which typically are relevant over a scale of many minutes to hours (3).

**Figure 1:**
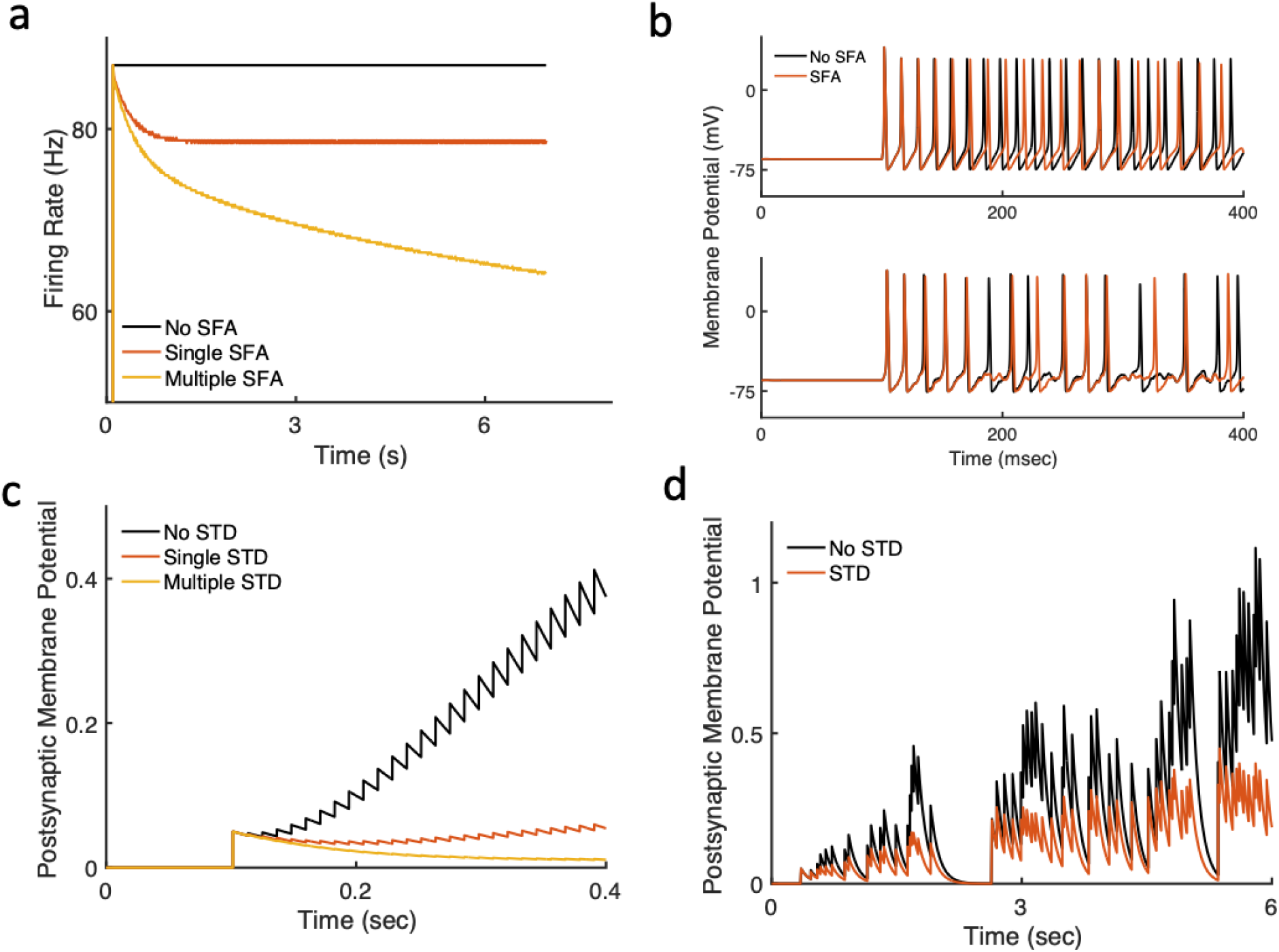
Spike frequency adaptation (SFA) and short-term synaptic depression (STD) are forms of negative feedback adaptation or gain control. (a) SFA leads to a decrease in action potential firing rate in response to a continuous stimulus. Examples show single and multiple timescale adaption. (b) Fewer action potentials are generated in noiseless (upper panel) and noisy (lower panel) stimulus conditions. STD decreases postsynaptic membrane potentials in response to (c) fixed presynaptic firing rates and (d) noisy presynaptic firing rates. Results are from Hodgkin-Huxley neuron models and synaptic-depression models with single or multiple timescales of adaptation (see Methods).

Despite the prevalence of negative feedback mechanisms, their effects on the properties of neural circuits are not clearly understood, especially as it relates to response timescales. Here, we define a neural model with pre- and postsynaptic multiple timescale adaptation and describe several emerging adaptation-related neural circuit properties at the level of local field potentials. We consider model results in the context of clinically recorded human neural activity, which has the potential to bridge local field potentials and human responses (10). Neural mechanisms of adaptation may be evident at the level of macroscale electrophysiological recordings and provide a window to underlying neural dynamics and overall circuit excitability.

### Neural Adaptation

First, we want to define a general model with which to examine multiple timescale adaptation. We define neural adaptation as processes that emphasize stimulus changes with computational properties similar to Spike Frequency Adaptation (SFA) and synaptic depression (STD) (11,12), which are common forms of negative feedback adaptation. SFA was first described over a century ago by Lord Adrian, who won a Nobel Prize for his discoveries (13). SFA is a type of high-pass filter or, equivalently, differentiator that emphasizes stimulus change. SFA linearizes the firing rate-input (f-I) curve of neurons (14) and can be approximated as (see also Supplemental Materials):

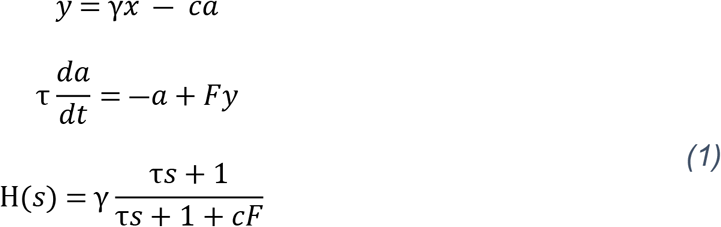

where *y* is the firing rate output, *x* is the mean current input, *a* is the adaptation variable, *c* and γ are gain constants, *F* governs the amount of adaptation (*0 < F < 1*), and *τ* is the effective adaptation time constant (15,16), which is smaller than the time constant of the underlying mechanism (17).

Short-term synaptic depression has input-output properties similar to SFA (11).

Specifically, synaptic depression can be modeled with the following nonlinear system of equations:

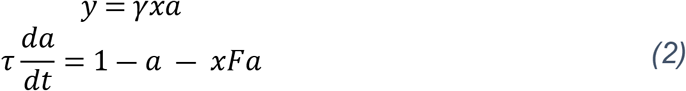

Due to the nonlinearity of the input *x* and adaptation variable *a* in both equations, we assume small fluctuations around *a*_0_ and *x*_0_ and linearize, which yields the following state space and transfer function equations (see Supplemental Materials):

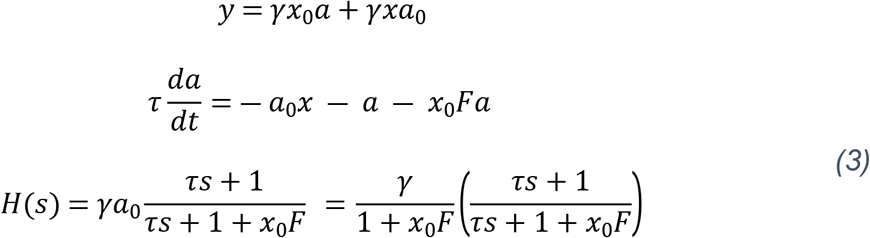

where *a* and *x* now represent small differences from steady-state values *a*_0_ and *x*_0_. The equations for SFA and synaptic depression are similar. However, in contrast to SFA, the non-linearity of synaptic depression leads to a gain factor that depends inversely on the input *x*_0_ since *a*_0_ = 1/(1 + *x*_0_ *F*), which has important consequences.

SFA and STD are not the only negative feedback mechanisms in neural circuits. Other negative feedback mechanisms have similar forms, including circuit delays, which can have transfer functions of the form *s/(τs + 1)*. Parallel feedforward networks in which inhibitory inputs lag behind excitatory inputs also lead to temporal differentiation with the same mathematical form (12). The core feature of these mechanisms is that the inhibitory feedback components lag excitatory components. The result is a derivative-like computation that emphasizes stimulus change and leads to decreasing temporal responsiveness to constant stimuli. When mechanisms such as SFA and STD occur in the same neural circuit, their differentiating properties can be additive (11,18), as expected with linear computations. Generally, multiple adaptive mechanisms, which are high-pass filters, exert negative feedback control, differentiate the input, and emphasize stimulus change. As we will see, multiple timescales of adaptive mechanisms can also give rise to history dependence and preserve information about past stimuli.

### Understanding Multiple Timescale Adaptation as Fractional Differentiation

Each differentiating process is described by a single-timescale adaptive process governed by an effective time constant *τ* However, each biological instantiation of negative feedback typically has a different timescale such that adaptation effectively has multiple timescales. We do not expect that each timescale will be the same or even similar. In fact, these timescales vary over at least an order of magnitude, such as demonstrated by mechanisms contributing to SFA (19) and STD (6). Then, how might we understand the computational effects of multiple timescales? We show that multiple timescales of adaptation are related to fractional dynamics, power laws, timescale invariance, and history dependence.

While single timescale processes may lie at one end of the timescale continuum, power law processes can be seen as lying at the other. Power laws can be thought of as including an infinite number of timescales such that they are scale invariant and no single timescale is preferred. Power law responses and multiple timescales of adaptation have long been observed in sensory processes (4,5) and may improve coding efficiency (20,21). The relationship between power laws and multiple timescales can be seen by arranging the definition of the gamma function with *x*=*λt* (5):

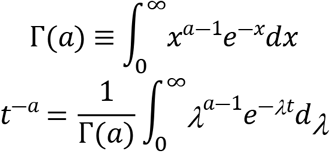

The integrand is comprised of weights *λ*^*a*-1^ for the exponential decay processes governed by timescale *λ*. In other words, power law decay can be thought of as composed of many exponential decay processes with properly balanced and varying timescales 1/*λ*. As described as early as Lord Adrian (13), decay processes are an example of neural adaptation in response to a step stimulus, or Heaviside step function. Practically, when these timescales occur in the internal variable of the feedback mechanism (e.g. Eq (1)), as few as three timescales can approximate a scale-invariant power law process over at least an order of magnitude (16,18). It is helpful to then notice that if instead we arranged the gamma function with *x*=*st*, integrate with respect to *t* and let *a*=-*α*, we obtain:

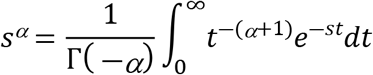

where it is immediately evident that *s^α^* is the Laplace transform of *t^(α+1)^*. We see that the power of *s* is related to the power law exponent. Regarding adaptation, we can see that *t^α^* is the output to a step stimulus, or Heaviside function, which is the integral of the delta function. We recall that *sX(s)* in the Laplace domain is the first derivative of *x(t)*, and that the delta function is the first derivative of the Heaviside function. We can then see that *s^α^*, which is a derivative of order *α* (which may be a non-integer), has power law dynamics in the time domain (8). In the time domain, the fractional derivative *D^α^* with order *α* can be written as (9,22):

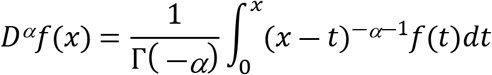

where the power law kernel on the righthand side of the equation is evident. Thus, at least if properly weighted, multiple timescale adaptation is intimately related to power law dynamics and can be succinctly characterized by fractional differentiation of order *α* for 0>*α* >1. The limits of integration from *0* to *x* demonstrate the history dependence of fractional differentiation.

When *α* = 1, the system performs a first derivative on the input, as described previously (11,12), and the mathematical computation is local in time. However, when *0* < *α* < 1 multiple timescales are present, stimulus history is maintained in the output, and adaptation is fundamentally non-local (Figure 2).

**Figure 2:**
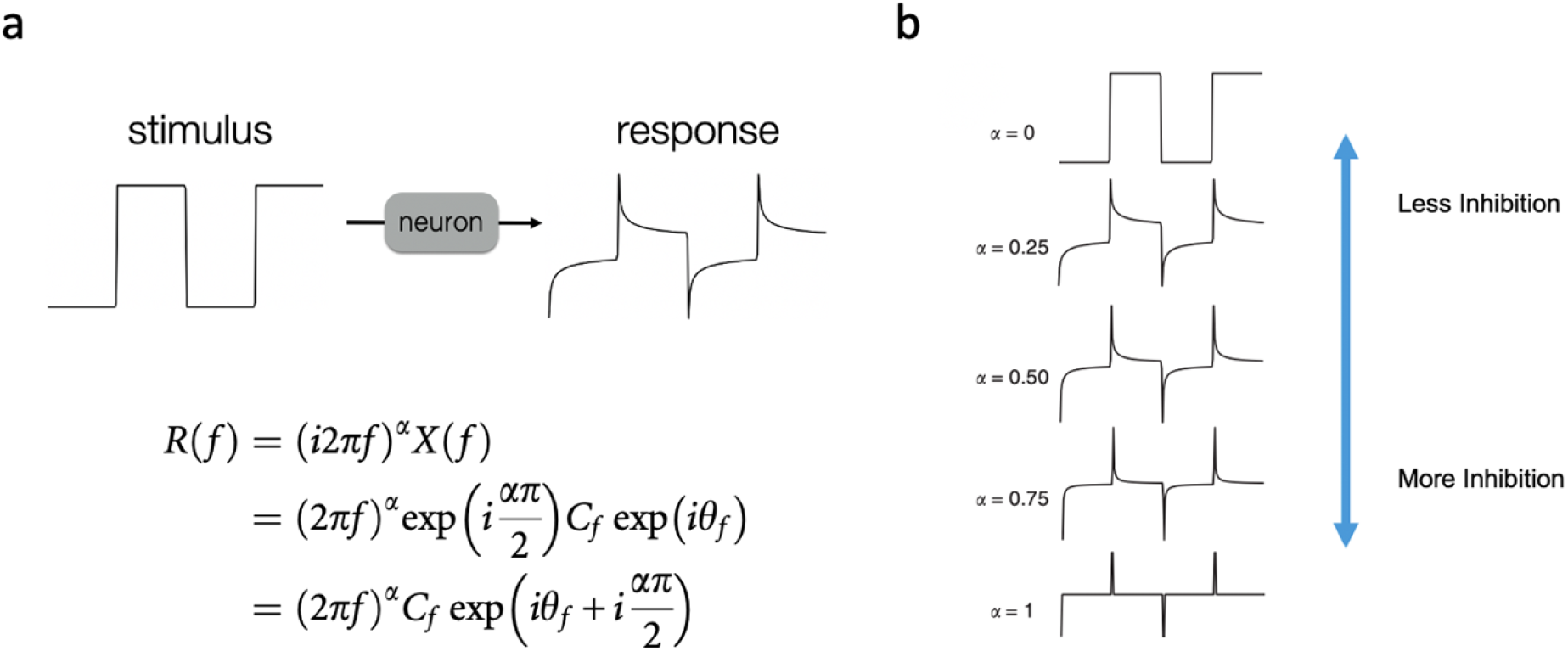
Neuronal spike frequency adaptation (a) approximates fractional differentiation with order ~0.1-0.5. Properties for the fractional derivative transfer function *(i2π)^α^* include a power law amplitude response and frequency-independent phase response (see also Supplemental Materials). (b) Higher fractional orders correspond to increased spike frequency adaptation, i.e., increased negative feedback inhibition.

### Adaptation, Power Spectra, and Broadband Power

Our previous work demonstrated multiple timescale adaptation consistent with fractional dynamics for *in vitro* and *in vivo* preparations. Neocortical neurons in slice preparations (8) and *in vivo* rodent thalamus and neocortical neurons (18) adapt to inputs in a manner consistent with fractional differentiation of order ~0.1-0.5. Here, we explore how these single neuron and neural circuit properties affect macroscale recordings such as EEG-recorded power spectral densities (PSD) and whether there is a PSD signature that may tell us something about underlying excitability.

Power spectra from local field potentials (LFPs) and EEG are generally thought to be dominated by post-synaptic membrane potentials (23), and here we consider summed post-synaptic membrane potentials as the source of the LFP and EEG signal. We focus on frequencies less than ~100 Hz and consider recorded LFP and EEG activity as resulting from a combination of high-pass and low-pass filtering mechanisms. Equations *(1)* and *(3)* describe the high-pass components within the transfer function, which provide negative feedback and define the adaptive timescales, and can be generally represented by:

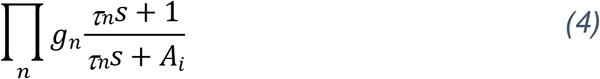

where the *n*th adaptive mechanism is represented by time constant *τ_n_* with associated gain constants *g_n_* and *A_n_*, where *A>1* (closer to *1* signifies less adaptation). We assume adaptive mechanisms are in series (see Supplemental Materials). To illustrate the characteristics of these adaptive mechanisms, we show the step response for each of three adaptative mechanisms, where the decay of the step response represents the timescale of adaptation (Figure 3). In this case, the time constants are 0.05, 0.5 and 5 sec, corresponding to three timescales. For comparison, we also plotted an approximate decaying power law, which inherently contains multiple timescales and is scale-invariant.

**Figure 3:**
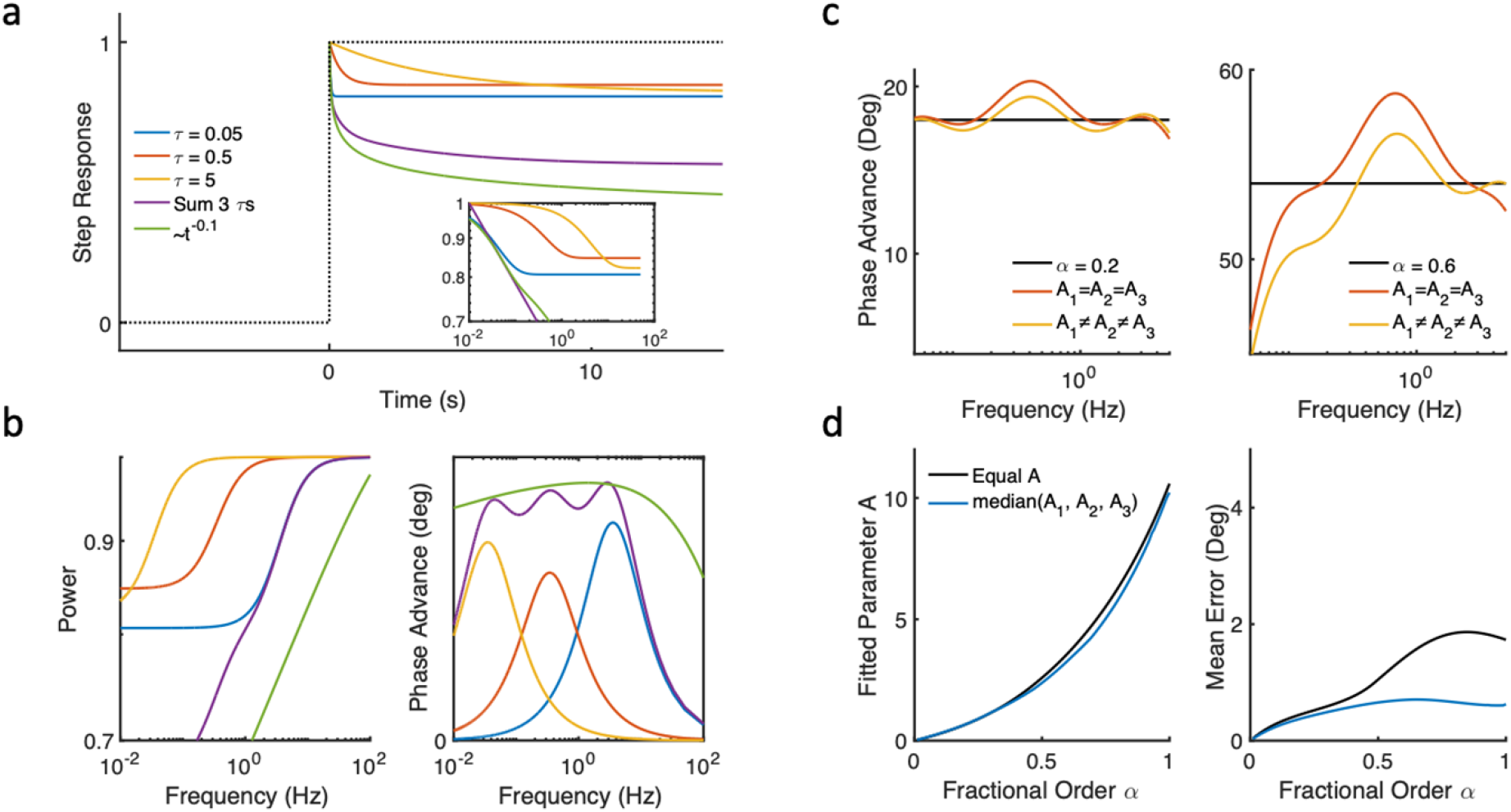
Multiple timescale adaptation can approximate fractional dynamics. (a) Step responses and (b) frequency domain characteristics of three high-pass filters with three timescales (*τ* = 0.05s, 0.5s, 5s). The sum of the three high-pass filters is similar to an approximate power law with exponent −0.1. (c) Fractional dynamics demonstrate a frequency-independent phase advance that can be approximated by multiple timescale adaptation with either one gain parameter *(A_1_*= *A_2_*= *A_3_*) or three gain parameters *(A_1_*≠ *A_2_*≠ *A_3_*). (d) Weighting each timescale equally (ie with one gain parameter) can provide a reasonable approximation of fractional dynamics, especially for α<0.5. The value of *A* (for one gain parameter) or median value of *A* (for three gain parameters) are similar as fractional order α increases.

Eq (4) may be constrained to ensure the multiple timescales are balanced such that there is not a single dominant timescale, as in the case of power laws or fractional dynamics (8). In other words, for a given set of *τ_n_*, balanced gain constants *A_n_* yield an approximate power law response. One approach involves fitting *A_n_* such that a frequency-independent phase results (16). For example, a system comprised of three logarithmically spaced time constants and equal *A* shows an approximate frequency independent phase advance (phase advances are additive), which is consistent with scale-invariant power law dynamics for frequencies ~0.01-10 Hz (Figure 3b, right panel).

We more rigorously examine how well three adaptive timescales can approximate fractional dynamics. Fractional dynamics implies a constant phase over some frequency range (Fig 2a, Supplemental Materials). Here, we use three timescales (0.05 s, 0.5 s, 5 s) to approximate the constant phase over the frequency range ~0.05-5 Hz. Examples are shown in Fig 2c under two conditions: with the same weight (*A_n_*) for each time scale and with separate weights. We minimize the difference between Eq. (4) and the constant phase defined by *α* using multidimensional unconstrained nonlinear minimization (Nelder-Mead). We find that equal weights for distinct well-separated timescales can approximate fractional dynamics, especially for *α* < 0.5 (Fig 2d).

Adaptative negative feedback mechanisms can be combined with a low-pass filter to represent a basic unit of neural computation. This fundamental computational neural unit can be thought of as spike generation through to postsynaptic membrane potentials, which is largely the source of LFP or EEG signals. The input is comprised of colored noise arriving at the neuron cell body, which can approximate neural activity and synchrony (24–28). This input causes action potentials that propagate along the axon and eventually give rise to postsynaptic potentials (PSPs). Action potentials and excitatory PSPs are high-pass filtered by adaptive mechanisms, while resistive and capacitive membrane characteristics serve as a low-pass filters (Figure 4).

**Figure 4:**
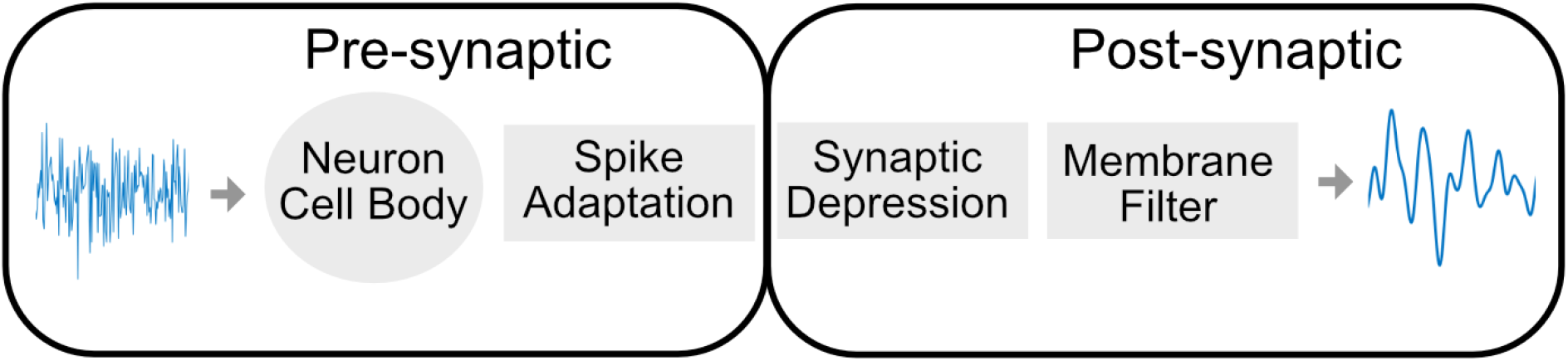
Model of neural components leading to LFP or EEG signals. Colored noise inputs are filtered by high-pass and low-pass filtering mechanisms. Neural circuit topologies with parallel feed-forward connections and asymmetric time delays have similar computational components.

The general transfer function *H(s)* for this series of low-pass and high-pass filters can be represented by:

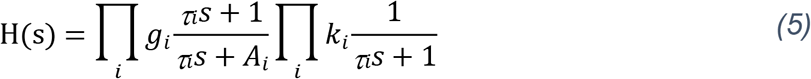

where *k_i_* are gain constants for the low-pass filters. This is a general form of a multiple timescale adaptive neural circuit element. When the high-pass filters include multiple timescales that are appropriately weighted, as previously discussed, the transfer function is approximately:

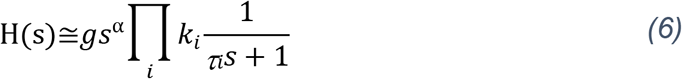

where *α* succinctly describes multiple timescale negative feedback of the system. The frequency-independent phase related to α can be used to find *A_i_* for a given set of *τ_i_* such that fractional dynamics are approximated (16) (Supplemental Materials). If Eq. (4) is being used to describe a neuron with multiple adaptation currents, then *A_i_*/*τ_i_* is approximately proportional to the adaptation channel conductances; a slightly modified form of the filter can yield a more precise correspondence between channel conductances and *A_i_* (16).

### Effects of adaptation on power spectral densities (PSD)

Negative feedback adaptation such as SFA and STD are typically relevant for frequencies less than 100 Hz since the fastest relevant time constants are ~10 msec or larger (19). The effect of multiple timescales is multiplicative in the frequency domain such that the largest effect of adaptation will be for lower frequencies. Thus, multiple timescale adaptation can change the PSD slope, as seen by *α* in Equation (6). To help exemplify this, we show the magnitude of the transfer function for a model with a single timescale of SFA, a single timescale of STD, and a low-pass filter:

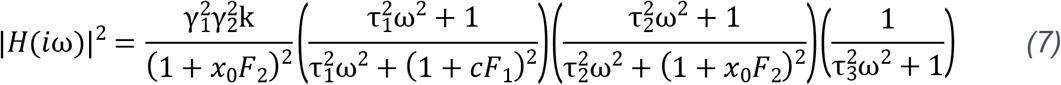

For SFA and STD, the denominator contains a frequency dependent term *τ^2^ω^2^* and a frequency-independent term, which is either *(1+cF)^2^* or *(1+x_0_F)^2^* for SFA and STD, respectively; all constants are positive. This form leads to sigmoid-like high-pass filters that are multiplied in the frequency domain with the low-pass filter and more strongly affect lower frequencies (Figure 5a).

**Figure 5:**
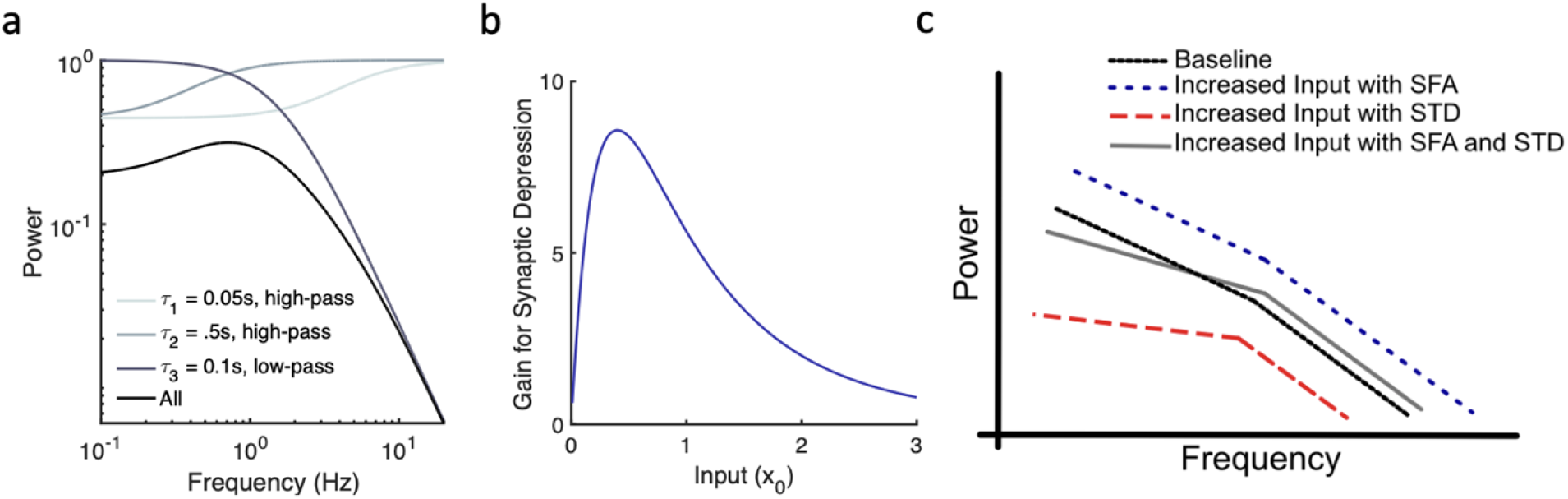
Adaptation affects the gain and slope of neural power spectral densities. (a) Example showing interaction of high-pass filters with low-pass filters. Eq (7) is plotted with three timescales (*γ*=*k*=*c*=*x_0_*=*1*, *F*=*0.5*). (b) Gain for synaptic depression initially increases and then decreases as *x_0_* increases (modeled via Eq (8) with *n*=*3*, *F*=*0.5*, *γ*=*2*) (c) Adaptation alters the slope of PSDs, decreases the overall gain, and affects lower frequencies more than higher frequencies. Adaptation interacts with input levels and includes Spike Frequency Adaptation (SFA) and Short-term Synaptic Depression (STD).

When there are multiple timescales of adaptation, the high-pass filters can be simplified such that for low to midrange frequencies (where adaptation is relevant) the magnitude of the transfer function is:

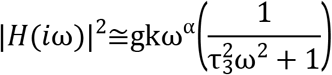

For high frequencies (eg >10-100 Hz), the PSD is that of the low-pass filter. For low to midrange frequencies, the slope of the PSD is approximately −*(2-a)*.

It is typically assumed that overall levels of power generally reflect overall levels of neural activity. Consistent with this, evidence from human single unit recordings suggests that, at least under some conditions, broadband increases in the PSD are correlated with increased neural firing rates (29). We expect this for SFA since Eq (1) is linear; if input *x* doubles, then the overall broadband power would double.

However, the non-linearity inherent in synaptic depression, as seen in Eq (3), means that increasing the input may lead to decreasing gain and overall power. This dependence results from the fundamental non-linearity of synaptic depression, where the amplitude of post-synaptic potentials depends on the product of pre-synaptic spikes and the amount of depression (6,12). The synaptic depression gain *g* = *γa_0_* has an inverse dependence on the input *x_0_*, since *a_0_*=*1/1+Fx_0_*. This means that for increasing levels of static (ie not time-varying) inputs, the output gain may decrease. Specifically, assuming that *F* is the same for all timescales (*0<F<1*), we see that the overall gain seen on a PSD for *n* timescales of synaptic depression would be:

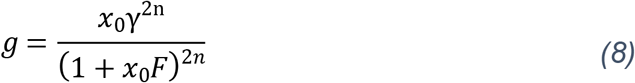

which means that as *x_0_* increases the gain increases until *dg/dx=0* at *x_0_ = 1/(2nF-F)*, after which further increases in *x_0_* cause the gain to decrease (Fig 5b). This non-linearity also means that the effect of STD depends on its input, which is in part determined by SFA. In other words, the input *x_0_* for STD is a function of SFA. Specifically, for a given input level, increasing SFA leads to fewer action potentials per second, which then decreases the input for STD. This decreases *a* in Eq (2) and results in decreased overall power in the postsynaptic membrane potential, especially for higher frequencies (since STD has a stronger effect on lower frequencies). This interaction between increased input levels, SFA, and STD, can lead to PSDs that show increased power for high frequencies and decreased power for low frequencies.

Several basic properties emerge from the general filter models in Eq (5):

1. Adaptation affects power for lower frequencies more than higher frequencies.
2. Increased input levels may decrease broadband power if synaptic depression is present.
3. Multiple timescale adaptation flattens the slope of power spectra according to *a*
4. The interaction of increased input and adaptation can decrease low frequency power in the output while increasing high frequency power.

These properties are schematically demonstrated in Figure 5c. To summarize, adaptation affects low frequencies due to its high-pass characteristics. Multiple adaptation timescales decrease power at lower frequencies, thereby flattening PSDs. The non-linearity of STD means that higher levels of steady-state input paradoxically lead to decreased broadband power. When only synaptic depression is present, increased input to the computational unit effectively increases the amount of negative feedback or adaptation in the neural circuity, which decreases power especially for lower frequencies. However, when SFA and STD are both present, moderately increased inputs decrease power for low frequencies and increase power for high frequencies, leading to a crossing of the baseline and final PSDs. The key difference between SFA and STD, in this respect, is the dependence on the input *x_0_* in the denominator of STD, which is the result of inherent non-linearities in STD that are not present in SFA. If inputs are further increased, then power for all frequencies may be decreased.

### Effects of multiple timescale feedback on power spectra

We want to examine how these properties may relate to PSDs from conductance-based model with spiking neurons and human EEG data from epilepsy patients. Previous work shows that the power spectra of EEG data demonstrate *1/f^β^* scaling (30,31), where *β* defines the exponent of the scaling and varies between ~2 and ~4. The shape or exponent of the power spectra may relate to the decay of synaptic inputs (30), ion channel or synaptic noise sources (32), or transitions between Up and Down states (33). Decreases in the exponent, ie a counter-clockwise PSD tilt, have been associated with visuomotor task-related neuronal activation, with an average intersection or pivot point of 28 Hz (34).

We see a similar PSD tilt when comparing previous data recorded near the seizure onset zone (SOZ) with data far from the SOZ for 91 patients implanted with invasive EEG electrodes (35,36). Specifically, power for frequencies less than ~1 Hz was decreased, while power for frequencies greater than ~1 Hz was increased near the SOZ compared to the non-SOZ (Figure 6). The cause of this tilt or crossing of the PSDs is not known, although the SOZ is generally considered to be hyperexcitable cortex.

**Figure 6:**
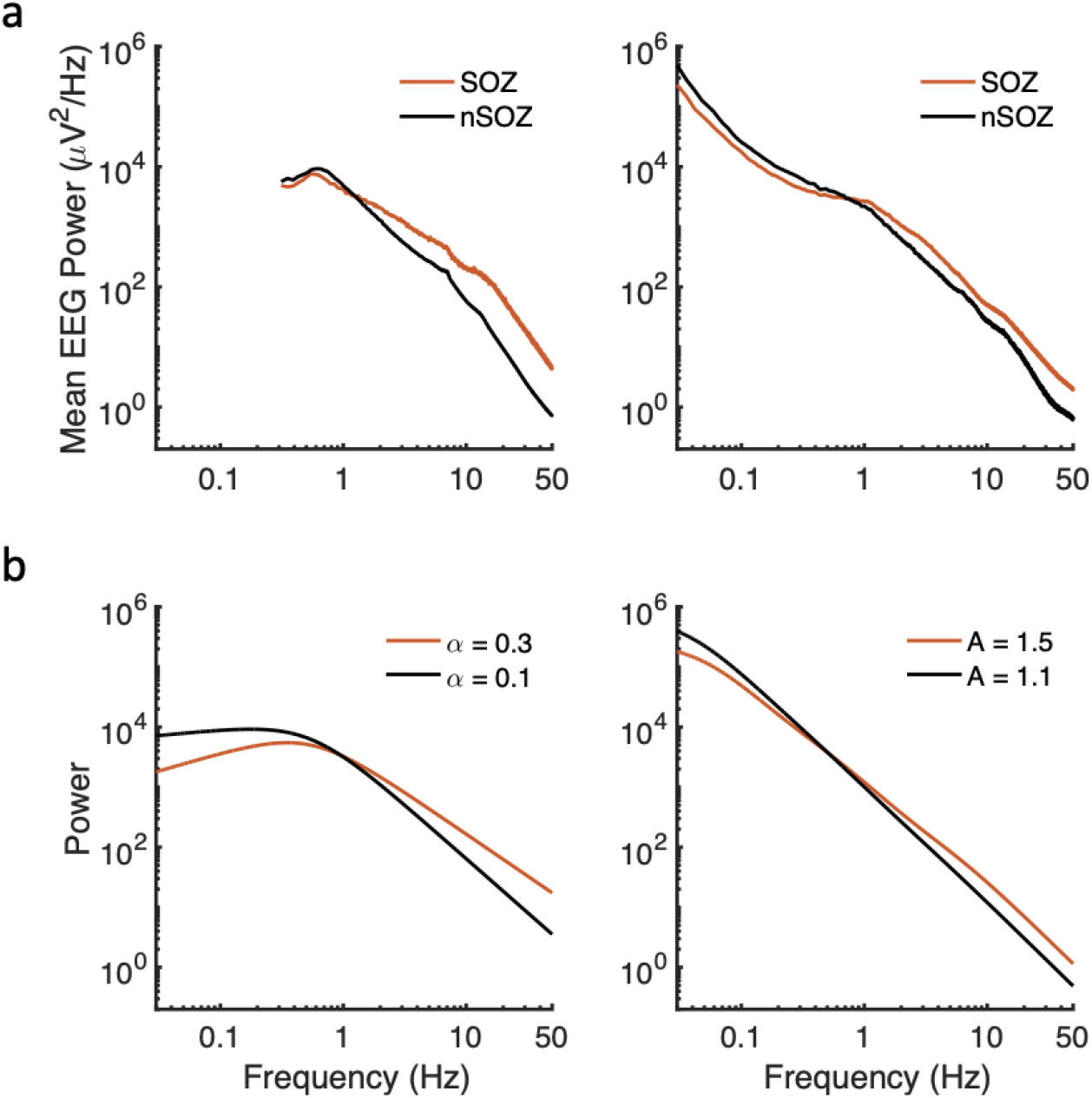
Power spectra changes related to the seizure onset zone and increased adaptation. (a) Previously published invasive EEG power spectra from 8 patients (left panel) (35) and from 83 patients (right panel) (36) showing decreased low frequency activity and increased high frequency activity near the seizure onset zone (SOZ) compared to the non-SOZ in epilepsy patients. (b) Similar shapes can be obtained from constant input models (with one low-pass timescale) when more negative feedback is present (e.g., increased *α* or *A*), implemented either via fractional dynamics (left panel; *α =0.3, g=0.7 or α=0.1, g=1;* for both *k=100, τ_m_=0.3 s)* or multiple timescales (right panel; *A_1_=A_2_=A_3_=1.1, g_1_=g_2_=g_3_=1.3* or *A_1_=A_2_=A_3_=1.5, g_1_=g_2_=g_3_=1.5;* for both *τ_1_=0.05 s, τ_2_=0.5 τ_3_=5 s, k=500, τ_m_=5 s*).

A tilt in the PSD is also seen for Eq (5) and Eq (6) when the amount of adaptation or negative feedback is altered (Figure 6b). Specifically, an increased *α* or *A* reflects increased negative feedback. Figure 6 shows that the power law exponent in the power spectral densities decreases as adaptation (negative feedback) increases. These examples don’t account for differing levels of input. To more realistically model PSD changes, we consider conductance-based models.

We implemented a feedforward neural network (Figure 7a) with conductance-based neurons (*n = 100*) that included three timescales of slow potassium currents, which implement spike frequency adaptation (16), as well as synaptic depression with two timescales (18). As predicted, increasing the input amplitude leads to decreased broadband power when STD is present. If the input is increased and STD is decreased, either directly or indirectly by increasing SFA, the resulting PSDs show a “tilt” (Figure 7b). Generally, filters that act directly upon the membrane current, like STD, more strongly effect resulting timescales than filters that act upon hidden variables, like SFA (37).

**Figure 7:**
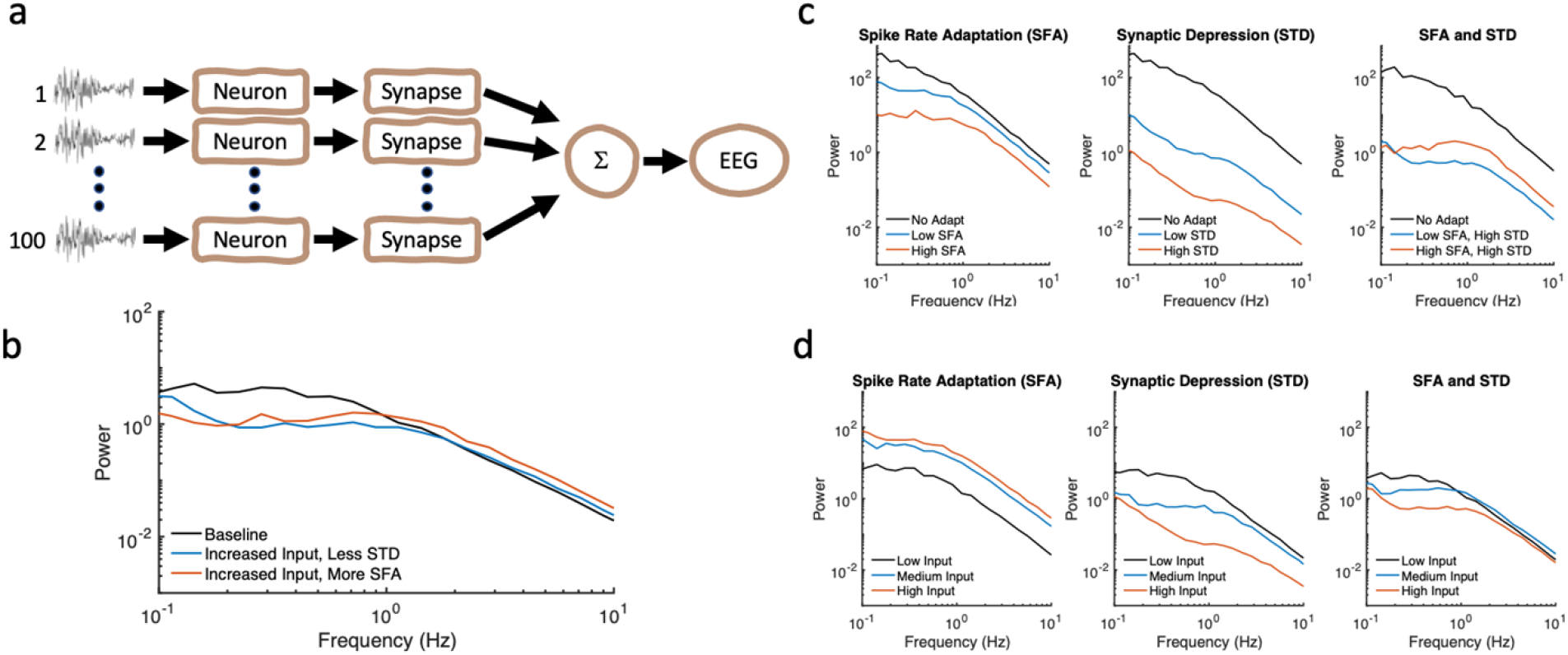
Spiking neuron network model with spike frequency adaptation (SFA) and synaptic depression (STD). (a) Feedforward neural network includes 100 conductancebased neurons with multiple timescales of SFA and STD. (b) PSDs show degrees of “tilt” or crossing when input is increased, and STD is decreased. Increasing SFA effectively decreases STD. (c) Increased adaptation, whether SFA or STD, generally leads to decreased power. However, increased SFA leads to decreased STD (rightmost panel). (d) Increasing input increases overall power when SFA is present. However, when STD is present, increasing the input leads to decreased overall power. When both SFA and STD are present, nonlinear effects are evident.

As expected, increased adaptation results in decreased power (Figure 7c). However, increasing SFA can increase broadband power if STD is present, as the input to STD is then decreased (Figure 7c, rightmost panel). Increasing the level of input to the neural circuit can have contrasting effects. If only SFA is present, then power increases. However, if only STD is present, then power in fact can decrease (Figure 7d), as above. When both SFA and STD are present, these two negative feedback mechanisms can have opposing effects, with STD having a greater effect on lower frequencies. Overall, the results from the conductance-based modeling are consistent with results from the filter models.

## Discussion

Negative feedback mechanisms, such as spike frequency adaptation (SFA) and shortterm synaptic depression (STD), are thought to be prevalent in human neural circuitry and yet their effects on local field potentials and EEG remain unclear. Here, we develop an approach for understanding multiple timescales of adaptation. This approach incorporates fractional dynamics, a form of non-integer order calculus that is related to power laws and has history dependence as a key feature. We find that alterations in neural excitability, consistent with mechanisms of SFA and STD, are most evident in low frequency (eg <10 Hz) EEG changes. In general, increased adaptation can lead to a counterclockwise tilt of the PSD. However, with concomitant increase of input, decreased adaptation can also lessen low frequency (<1 Hz) activity. Counterintuitively, we find that increased inputs can decrease broadband power via the effects of STD.

Typically, the SOZ is considered hyperexcitable cortex. Adaptive mechanisms may play a role in dampening hyperexcitability in SOZ during the interictal period and fail during seizure initiation (38,39). Previous interictal results show decreased low frequency (<1 Hz) and increase high frequency (2-50 Hz) activity near the SOZ (35,36). Present results suggest that that this pattern is consistent with an interaction between altered adaptation and increased input.

Previously, we have described computational properties consistent with fractional-order derivatives in rodents. Single neocortical neurons from brain slices were found to adapt with multiple timescales in response to injected currents of varying amplitudes and variances (8). When compared with a time-varying input, the firing rate output was a waveform consistent with the fractional derivative of the input. The order of the fractional derivative was found to be approximately 0.15, and this order was increased or decreased when slow after-hyperpolarization (sAHP) currents were increased or decreased, respectively. Similar results were obtained from *in vivo* rodent neurons (18). Adaptation to whisker movements of anesthetized rats was found in thalamic and neocortical neurons with orders of approximately 0.2 and 0.5, respectively. This is consistent with the idea that multiple SFA and STD processes along the sensory pathway contribute to the adaptation seen in the neocortex. We have previously suggested an approach for fitting multiple exponential adaptation processes to achieve a desired order of fractional differentiation, which approximates a power law (16), thus relating adaptation mechanisms to neural circuit responses.

Fractional differentiation and fractional derivatives are fundamentally related to power laws (5,9,22,40). These mechanisms may promote efficient encoding and information transmission (4,20,21). In general, adaptation has been linked to prediction (11) and memory (41), and multiple timescales may be critical for performing computational tasks (42). As we have seen, mechanisms of pre- and post-synaptic adaptation can interact, and it is critical to consider them together (7).

Here, we presented a neural circuit model with SFA and STD to relate neuron-level adaptation mechanisms with changes in human EEG. These results suggest that EEG changes in the frequency range ~0.01-1 Hz may reflect, at least in part, changes in the negative feedback of neural circuitry. Analysis of this EEG frequency range may provide estimates of neural excitability. Neural adaptation may be evident in macroscale electrophysiological recordings and provide a window to understanding neural circuit excitability.

## Methods

### Biophysical modeling

The neural network consisted of Hodgkin-Huxley (HH) neurons (2,43) and synapses that incorporated depression (6). The single-compartment, conductance-based Hodgkin-Huxley (HH) model neuron was used with standard parameters: *G*_Na_ = 120, *G*_K_ = 36, and *G*_Leak_ = 0.3 mS/cm^2^; *E*_Na_ = 50, *E*_K_ = −77, and *E*_Leak_ = −54.4 mV; and *C = 1 μF/cm^2^*. Equations were solved numerically using fourth-order Runge-Kutta integration with a fixed time step of 0.05 msec, and spike times were identified as the upward crossing of the voltage trace at −10 mV (resting potential = −65 mV) separated by more than 2 msec. Input stimuli were zero mean exponentially-filtered (*τ* = 1 msec) Gaussian white noise stimulus with SD that was 10, 15, or 20 μA/cm^2^, which was used for the Low Input, Medium Input, and High Input conditions. Without adaptation, these inputs corresponded to ~1, 10, and 20 Hz spike output.

### Spike Frequency Adaptation

Three slow adaptation currents were added to the HH neurons as previously (8). For each AHP current, an additional current, or term, was added to the equation for *dV/dt* of the form –*G*_AHP_*a*(*V* – *E*_reversal_), where *a* was incremented by one after each spike and decayed according to *da/dt = −a/τ*, with *τ* = [0.3, 1, 6] sec and *G*_AHP_ = [0.05, 0.006, 0.004]*G*_Leak_. Low SFA employed *G*_AHP_, while High SFA was 5*G*_AHP_.

### Synaptic Depression

Synaptic depression was modeled as previously (6,18), with facilitation omitted. Depression variables (*d* and *D*) in the synapses relaxed exponentially to one with time constants of 0.6 and 9 s for fast depression and slow depression, respectively. For depression each input spike decreased *d* and *D* to new values of 0.4*d* and 0.995*D*, respectively. These parameters are similar to those found in the best-fit model for short-term synaptic depression (6). This is equivalent to calculating the synaptic conductance amplitudes according to:

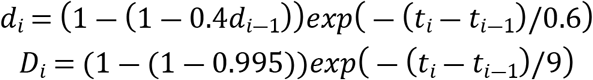

where *t_i_* is the *i*th presynaptic spike time in seconds. The synaptic conductance amplitude, *G* = *d D*, for each neuron was then summed and filtered by an exponential with a time constant of 0.3 s. Presynaptic amplitudes are summed across neurons, and Welch’s Method was used to find PSDs. The above parameters were used for the “High STD” condition. For Low STD, each input spike decreased *d* and *D* to new values of 0.7*d* and 0.999*D*.

